# The absence of gastrointestinal redox dyshomeostasis in the brain-first rat model of Parkinson’s disease induced by bilateral intrastriatal 6-hydroxydopamine

**DOI:** 10.1101/2022.08.22.504759

**Authors:** Jan Homolak, Mihovil Joja, Gracia Grabaric, Emiliano Schiatti, Davor Virag, Ana Babic Perhoc, Ana Knezovic, Jelena Osmanovic Barilar, Melita Salkovic-Petrisic

## Abstract

The gut-brain axis plays an important role in Parkinson’s disease (PD) by acting as a route for vagal propagation of aggregated α-synuclein in the gut-first endophenotype and as a mediator of gastrointestinal dyshomeostasis via the nigro-vagal pathway in the brain-first endophenotype of the disease. One important mechanism by which the gut-brain axis may promote PD is by regulating gastrointestinal redox homeostasis as overwhelming evidence suggests that oxidative stress plays a key role in the etiopathogenesis and progression of PD and the gastrointestinal tract maintains redox homeostasis of the organism by acting as a critical barrier to environmental and microbiological electrophilic challenges. The present aim was to utilize the bilateral intrastriatal 6-hydroxydopamine (6-OHDA) brain-first PD model to study the effects of isolated central pathology on redox homeostasis of the gastrointestinal tract. Three-month-old male Wistar rats were either not treated (intact controls; CTR) or treated bilaterally intrastriatally with vehicle (CIS) or 6-OHDA (6-OHDA). Motor deficits were assessed with the rotarod performance test and the duodenum, ileum, and colon were dissected for biochemical analyses 12 weeks after the treatment. Lipid peroxidation, total antioxidant capacity, low-molecular thiols, and protein sulfhydryls, the activity of total and Mn/Fe superoxide dismutases, and total and azide-insensitive catalase/peroxidase were measured. Univariate and multivariate models of redox biomarkers provide solid evidence against the existence of pronounced gastrointestinal redox dyshomeostasis. The results indicate that the dysfunction of the nigro-vagal system and not motor deficit may be a key mediator of gastrointestinal dyshomeostasis in brain-first 6-OHDA-induced rodent models of PD.

## Introduction

Parkinson’s disease (PD) is a chronic progressive neurodegenerative condition characterized by degeneration of dopaminergic neurons in substantia nigra (SN) pars compacta that results in the development of bradykinesia, tremor at rest, rigidity, and postural instability [1]. Although the etiopathogenesis of the disease remains to be elucidated accumulating evidence points to the involvement of the gastrointestinal tract as i) the prodromal non-motor symptoms affecting the gastrointestinal tract (e.g. dysphagia, delayed gastric emptying, and constipation) are prevalent and precede the motor phase of PD (sometimes by decades)[2–4]; ii) a stereotypical spreading pattern of α-synuclein pathology [5, 6] supports the hypothesis that misfolded α-synuclein may originate from the gut [7]; and iii) mechanistic animal studies clearly demonstrate that pathophysiological events in the gastrointestinal tract are sufficient to trigger and promote the development of the central nervous system (CNS) pathology resembling PD (e.g.[8–10]). Based on the aforementioned evidence a body-first hypothesis of PD has been proposed with the gastrointestinal tract considered as the most likely site of early molecular pathophysiological events [11].

In contrast, some studies suggest that in a considerable proportion of patients PD does not propagate in concordance with the Braak staging system [12, 13]. Furthermore, although highly prevalent, gastrointestinal symptoms are not present in all patients diagnosed with PD, and they do not always appear before the onset of motor symptoms [2, 14]. Consequently, it is evident that in some patients a brain-first hypothesis provides a more accurate explanation of the PD progression.

Based on the aforementioned data, a working model has been proposed which recognizes PD as a complex disease comprised of at least two clusters of phenotypes (brain-first and gut-first)[11, 15]. The gut-brain axis plays an important role in both subtypes acting as a route for vagal propagation of aggregated α-synuclein in the gut-first phenotype and as a mediator of gastrointestinal dyshomeostasis via the nigro-vagal pathway in the brain-first phenotype (e.g. [16]). Nevertheless, the mechanisms by which the gut-brain axis may contribute to the propagation of the disease and the appearance of gastrointestinal symptoms remain poorly understood and challenging to study due to overlapping brain and gut pathology in animal models.

In this context, the CNS-targeted 6-hydroxydopamine (6-OHDA) rodent models provide a unique way to study the effects of the brain-first predominant subtype of the disease on the pathophysiological alterations in the gut as the toxin cannot cross the blood-brain barrier. The central 6-OHDA administration model was first introduced by Ungerstedt following the idea that high selectivity of the toxin towards the dopamine uptake sites may result in a specific nigrostriatal dopaminergic lesion [17]. Some groups already utilized the model to study the mechanisms of gastrointestinal dyshomeostasis in the context of the brain-first PD-like nigrostriatal lesion primarily with the focus on gastrointestinal motility (e.g. [18–20]). Other pathophysiological alterations such as decreased expression of the occludin barrier proteins [21] and impaired production of mucus [22] have also been reported.

Interestingly, the effects of brain-first PD-like lesion have, to the best of our knowledge, never been explored in the context of intestinal redox homeostasis regardless of the fact that i) there is overwhelming evidence that redox dyshomeostasis and oxidative stress play a critical role in the pathophysiology of neurodegeneration [23] and the etiopathogenesis of PD [24, 25]; ii) The structure and function of the gastrointestinal barrier and the regulation of redox homeostasis are highly interdependent [26, 27]; and iii) the gastrointestinal tract is considered to be a „free radical time bomb“due to its constant and inevitable exposure to foreign substances and microorganisms with substantial electrophilic potential [28]. Considering that other neurotoxin-based „brain-first“models of neurodegeneration (e.g. the streptozotocin-induced rat model of sporadic Alzheimer’s disease [29] with some resemblance to the 6-OHDA models of PD [30]) demonstrate pronounced redox dyshomeostasis that may contribute to the progression of the disease phenotype [31, 32], we hypothesized that intrastriatal administration of 6-OHDA may produce similar alterations and promote systemic and neuro-inflammation in the rat model of PD contributing to the progression of neuropathology.

The present aim was to measure redox biomarkers along the gastrointestinal tract in a rat model of brain-first PD induced by bilateral intrastriatal administration of 6-OHDA.

## Materials and methods

### Animals

The experiment was performed on three-month-old male Wistar rats (N=37) from the Department of Pharmacology (University of Zagreb School of Medicine, Zagreb, Croatia). All procedures were approved by the University of Zagreb School of Medicine Ethics Committee and the Croatian Ministry of Agriculture (EP 186 /2018; 380-59-10106-18-111/173) and complied with current institutional, national, and international guidelines (The Animal Protection Act, NN102/17; NN 47/2011; Directive 2010/63/EU).

The rats were housed 2-3/cage with a 7 AM/7 PM 12-hour light cycle with controlled humidity (40-70%) and temperature (21-23°C). Tap water and standardized food pellets were available *ad libitum* and bedding was changed twice per week. Before the model induction, the animals were randomized into 3 groups – i) intact controls (CTR; N=9) that would not undergo any procedure (control for the effects of sham procedure); ii) control animals (CIS; N=14) that would receive a bilateral intrastriatal injection of vehicle (control for the effects of 6-OHDA); and iii) Parkinson’s disease model (6-OHDA; N=14) that would receive a bilateral intrastriatal injection of the 6-OHDA solution. The experimental design was unbalanced with respect to the number of animals assigned to each group due to a greater anticipated death rate in the experimental arms with invasive surgical procedures (i.e. CIS and 6-OHDA).

### Intrastriatal administration of 6-hydroxydopamine

6-OHDA was dissolved in 0.02% w/v ascorbic acid in sterile saline on the day of the procedure. The animals from the CIS and the 6-OHDA groups were anesthetized by intraperitoneal administration of ketamine (70 mg/kg) and xylazine (7mg/kg). The skin was surgically opened and the skull was trepanated with a drill. 2 μl of the 6-OHDA solution (4 μg 6-OHDA/μl; 6-OHDA) or an equal volume of vehicle (CIS) was administered into each hemisphere at stereotaxic coordinates: 0 mm posterior, 2.8 mm lateral, and 7 mm ventral relative to bregma and pia mater respectively [33]. The rats were monitored for 24 hours after the surgery.

### Rotarod performance test

The rotarod performance test was used to assess motor deficits 3 months after the model induction. Briefly, the animals are placed on an elevated cylinder rotating at a constant speed and the time-to-fall is recorded. Each animal is first trained to avoid the bias introduced by habituation to the task and learning, and each time-point consisted of equally spaced two consecutive trials with a 180 s cut-off time.

### Sample preparation

After 3 months of model induction, the animals were euthanized in deep anesthesia achieved by intraperitoneal administration of ketamine (70 mg/kg) and xylazine (7 mg/kg). Duodenum, ileum, and colon were excised and luminal contents were removed with a syringe filled with ice-cold phosphate-buffered saline (PBS). The samples were cut open, rinsed again in ice-cold PBS, snap frozen in liquid nitrogen, and stored at −80°C until further analyses. Duodenum was sampled 1 cm distal from the pylorus, ileum 1 cm proximal from the caecum, and colon at the midline between the caecum and the sigmoid. The samples were homogenized using an ultrasonic homogenizer (Microson Ultrasonic Cell 167 Disruptor XL, Misonix, Farmingdale, NY, SAD) in 7.5 pH lysis buffer containing 150 mM NaCl, 50 mM Tris-HCl pH 7.4, 1 mM EDTA, 1% Triton X-100, 1% sodium deoxycholate, 0.1% SDS, 1 mM PMSF, protease inhibitor cocktail (Sigma-Aldrich, Burlington, MA, USA) and phosphatase inhibitor (PhosSTOP, Roche, Basel, Switzerland). Homogenates were spun down for 10 minutes at 4°C and a relative centrifugal force of 12 879 g, and supernatants were stored at −80°C. Protein concentration (used as a covariate for other variables to account for differences in lysis efficacy) was measured using the Bradford reagent (Sigma-Aldrich, USA) and bovine serum albumin dissolved in the same lysis buffer for the generation of the calibration curve.

### Thiobarbituric acid reactive substances assay

Thiobarbituric acid reactive substances (TBARS) assay was used to quantify end products of lipid peroxidation as described by Prabhakar et al. [34] and modified in [35]. 12 μl of each sample was mixed with 120 μl of the TBA-TCA reagent (w/v 0.04% thiobarbituric acid (Kemika, Croatia) in 15% trichloroacetic acid (Sigma-Aldrich, USA). The samples were diluted with 70 μl of ddH_2_O, vortexed, and incubated in a heating block set at 95°C for 20 minutes in perforated microcentrifuge tubes. The reaction was monitored by visual inspection of the experimental and standard samples and the incubation time was prolonged if needed. The colored product was extracted in 220 μl of n-butanol. The absorbance of the butanol extract was measured at 540 nm in a 384-well plate using the Infinite F200 PRO multimodal plate reader (Tecan, Switzerland). The concentration of TBARS was estimated from the standard curve of the MDA tetrabutylammonium (Sigma-Aldrich, USA) processed in parallel. The extraction procedure was adapted if necessary by adjusting the sample input volume, increasing the duration of the heating step, and modifying the volume of n-butanol used in the extraction procedure based on the partitioning coefficient of the colored product. All adjustments to the procedure were made with standard samples processed in parallel to enable the comparison of standardized estimates.

### Ellman’s procedure for determination of low-molecular-weight thiols and protein sulfhydryls

Low molecular weight thiols (with glutathione (GSH) being the most abundant) and free protein sulfhydryls (SH) were quantified by measuring 5-thio-2-nitrobenzoic acid (TNB) following the reaction of 5,5⍰-dithio-bis(2-nitrobenzoic acid) (DTNB) with the supernatant and the pellet after precipitation of proteins with sulfosalicylic acid [31, 36, 37]. Briefly, 25 μL of each homogenate was incubated with 4% w/v sulfosalicylic acid in ddH_2_O for 1 h on ice in a 1:1 volumetric ratio. The samples were centrifuged for 10 min at 10 000 g and both the supernatant and the pellet were reacted with w/v 4 mg/ml DTNB in w/v 5% sodium citrate in ddH_2_O for 10 minutes. The absorbance of the supernatant from both reactions was read at 405 nm using the Infinite F200 PRO multimodal microplate reader (Tecan, Switzerland) and the concentration was calculated using a molar extinction coefficient of 14,150 M^−1^cm^−1^.

### Superoxide dismutase activity

The activity of superoxide dismutase (SOD) was determined indirectly by the inhibition of the 1,2,3-trihydroxybenzene (THB) autoxidation [38, 39] using a modified protocol [37, 40]. The homogenates (3 μl for duodenum or 6 μl for ileum and colon) were placed in a 96-well plate and incubated with a 100 μl of the buffer for measuring total SOD activity (80 μL of the THB solution (60 mM THB dissolved in 1 mM HCl) added to 4000 μL of 0.05 M Tris-HCl and 1 mM Na2EDTA (pH 8.2)) or Fe/Mn-SOD activity ((80 μL of the THB solution (60 mM THB dissolved in 1 mM HCl) added to 4000 μL of 0.05 M Tris-HCl, 1 mM Na2EDTA, 2 mM KCN (pH 8.2)). Absorbance increment at 450 nm reflecting THB autoxidation was measured using the Infinite F200 PRO multimodal microplate reader (Tecan, Switzerland) with 30 s kinetic interval time cycles for 300 s.

### Hydrogen peroxide dissociation rate

Sample-induced H_2_O_2_ dissociation was measured to assess the catalase and residual peroxidase activity using the method originally proposed by Hadwan [41] and modified in [42]. Duodenum (4 μl), ileum (7 μl), and colon (10 μl) samples were first incubated with 100 μl of the Co(NO_3_)_2_ working solution (5 mL of Co(NO_3_)_2_ × 6 H_2_O (0.2 g dissolved in 10 mL ddH_2_O) mixed with 5 mL of (NaPO_3_)_6_ (0.1 g dissolved in 10 mL ddH_2_O) added to 90 mL of NaHCO_3_ (9 g dissolved in 100 mL ddH_2_O)) followed by the H_2_O_2_ working solution (40 μl of 10 mM H_2_O_2_ in 1xPBS) to obtain baseline values for the adjustment of endogenous H_2_O_2_ and/or chemical interference. The same procedure was repeated with samples first incubated with the H_2_O_2_ working solution and the Co(NO_3_)_2_ working solution added at t_1_= 60 s to assess the dissociation rate. The H_2_O_2_ concentration in each well was determined from the oxidation rate of cobalt (II) to cobalt (III) in the presence of bicarbonate ions using the carbonato-cobaltate (III) complex ([Co(CO_3_)_3_]Co) absorbance peak at 450 nm with the Infinite F200 PRO multimodal microplate reader (Tecan, Switzerland). The same procedure was repeated with a modified H_2_O_2_ working solution containing 0.025 mM sodium azide (AZD) to inhibit catalase activity (and measure residual peroxidase dissociation rate)[43]. The concentration of H_2_O_2_ was estimated from the standard model obtained by reacting the Co(NO_3_)_2_ working solution with serial dilutions of H_2_O_2_ in 1xPBS. The reaction time and optimal volume of the sample and working solutions were determined based on pilot experiments to ensure optimal sensitivity of the assay.

### Nitrocellulose redox permanganometry

Nitrocellulose redox permanganometry (NRP)[44] was used to determine the total reductive/antioxidative capacity of intestinal samples. Briefly, 1 μl of the homogenate was pipetted onto the nitrocellulose membrane (Amersham Protran 0.45; GE Healthcare Life Sciences, Chicago, IL, USA) and left to dry out at room temperature. The membrane was immersed into the KMnO_4_ solution (0.2 g KMnO_4_ in 20 ml ddH_2_O) for 30 s, rinsed under running dH_2_O, and left to dry out at room temperature. The MnO_2_ precipitate trapped on the membrane was quantified in Fiji (NIH, Bethesda, MD, USA).

### Data analysis

Data were analyzed in R (4.1.3) in concordance with the guidelines for reporting the evidence from animal studies [45]. The experimenters were not blinded and the animals were assigned to experimental groups based on stratified randomization (in regards to body mass, home cage allocation, and litter). Survival analysis was done using survfit algorithm from the survival package [46] and the results were reported using survminer [47]. The animals were monitored daily since the model induction and all animals that reached the final time-point (85 days) were censored. The group was defined as a stratum. A similar approach was used for the analysis of the rotarod performance test to account for the pronounced ceiling effect as most animals from the control groups (CTR, CIS) reached the cut-off time of 180 s. The cut-off time was not modified during the experiment as the aim was to confirm that the pronounced motor deficit was present in the animals treated with 6-OHDA and the underestimated estimates already informed of a substantial biological effect. The performance was measured as the time spent on the rotating cylinder in two subsequent 180 s trials. Failure to reach the cumulative cut-off time was defined as an *event* and all animals that reached the cut-off time were censored. The α was set at 5%. Redox biomarkers were analyzed using linear regression with the variable of interest used as the dependent variable and group/treatment allocation used as independent variables. Protein concentration was introduced as a covariate in each model to adjust for potential differences introduced during the lysis procedure (i.e. loading control). For THB autoxidation, the difference between the final and baseline absorbance (δTHB) was used as the dependent variable and baseline absorbance was introduced as a covariate to account for potential chemical interference. For the H_2_O_2_ dissociation rate, the difference in the absorbance between the baseline and the final time-point was used as the dependent variable, and baseline sample absorbance (incubation with the Co(NO_3_)_2_ working solution before the addition of the H_2_O_2_ working solution) was defined as a covariate. Model assumptions were checked using visual inspection of residual and fitted value plots and transformations were used where appropriate. Model outputs were reported as point estimates with 95% confidence intervals, and differences between groups were reported as ratios of least square means with accompanying 95% confidence intervals. Multivariate exploration was done by the principal component analysis of centered and scaled variables. The results were presented as biplots with respective contributions of redox biomarkers to the first two components.

## Results

### Bilateral intrastriatal administration of 6-hydroxydopamine induces pronounced motor deficits confirmed by the rotarod performance test

As expected the 6-OHDA model was characterized by a relatively large drop-out rate throughout the experiment while no fatal outcomes were observed in either of the two control groups (CTR, CIS). The fatality rate associated with the 6-OHDA administration was most pronounced in the first month of the experiment (50%) and stabilized thereafter reaching 57% in the final time-point (Fig 1A). The rotarod performance test showed pronounced motor deficits in the 6-OHDA group and no difference between the intact and the vehicle-treated animals 3 months after the model induction (Fig 1B, C).

**Fig 1.**
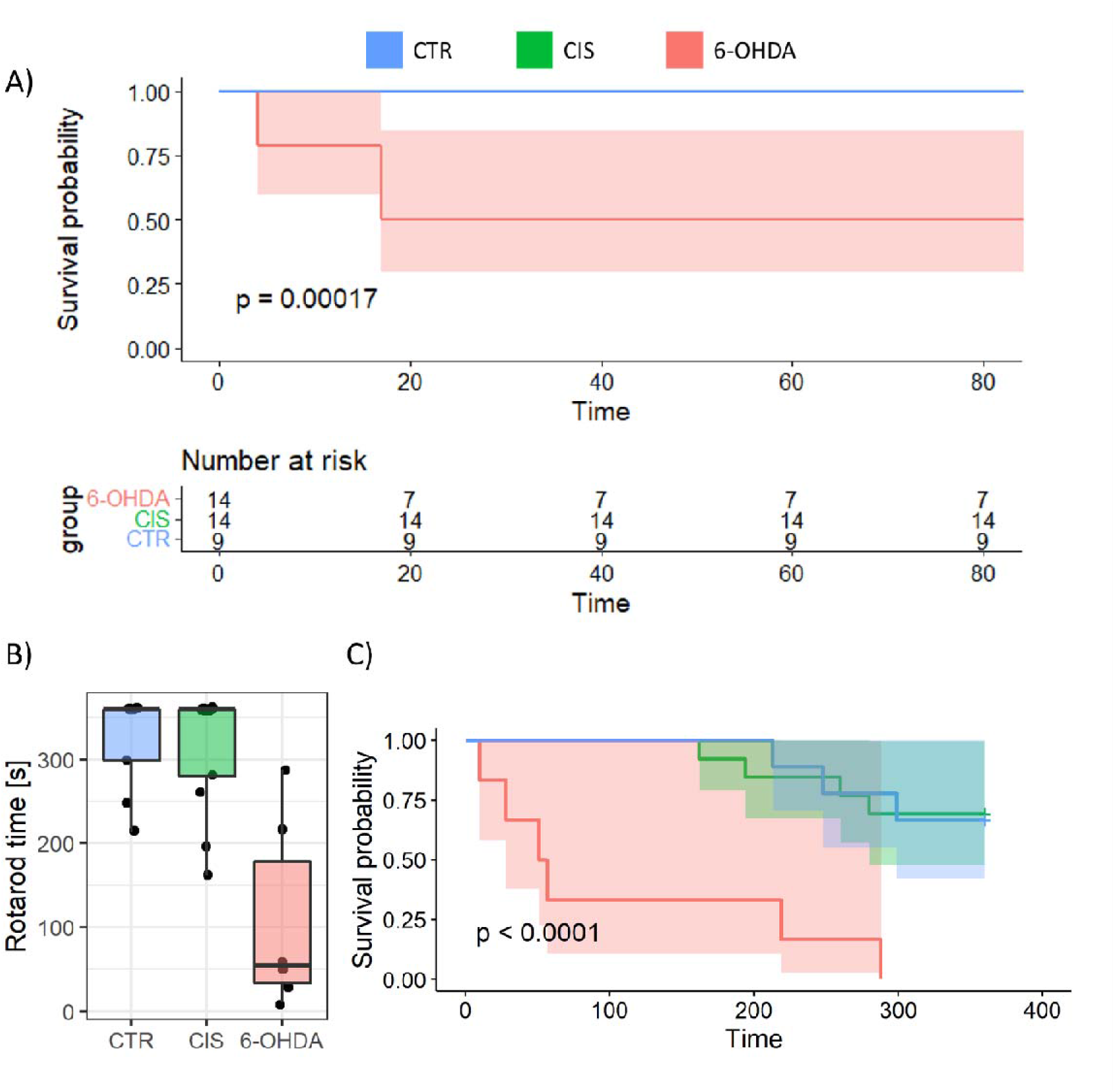
Survival and motor performance of 6-OHDA-treated and the control rats. A) Survival curve and the risk table for the experiment demonstrate a pronounced drop-out rate in the 6-OHDA group. B) Rotarod performance test 3 months after the model induction demonstrating substantial motor deficits in the 6-OHDA-treated rats. C) Survival analysis of the rotarod performance test to account for the pronounced ceiling effects in the control groups (CTR, CIS). Time represents cumulative rotarod performance time 3 months after the model induction. CTR – intact controls; CIS – control animals intrastriatally treated with the vehicle; 6-OHDA – rat model of Parkinson’s disease intrastriatally treated with 6-hydroxydopamine.

### Motor deficits induced by the bilateral intrastriatal 6-hydroxydopamine are not accompanied by intestinal redox dyshomeostasis

Analysis of redox homeostasis in 3 different segments of the intestine (duodenum, ileum, and colon) indicates that bilateral intrastriatal administration of 6-OHDA is not sufficient to perturb redox homeostasis of the gastrointestinal tract in the rat model of Parkinson’s disease 12 weeks after model induction. In contrast to what was expected, lipid peroxidation was decreased in the intrastriatally-treated animals (Table 1, Fig 2). Interestingly, the effect was most pronounced in the duodenum and more pronounced in the 6-OHDA-treated animals in comparison with sham-operated controls. Other redox biomarkers remained largely unchanged across treatments and intestinal segments indicating small effects and large uncertainty of estimates. The H_2_O_2_ dissociation rate demonstrated a similar pattern of reduced capacity in the 6-OHDA reversed in the presence of catalase inhibitor AZD both in the duodenum and in the ileum suggesting that the small intestine of the 6-OHDA-induced PD model may be more reliant on the activity of catalase for removal of H_2_O_2_. Nevertheless, the effects of 6-OHDA on both the total H_2_O_2_ dissociation rate and AZD-resistant H_2_O_2_ dissociation capacity were relatively small (especially considering that the activity of catalase frequently changes exponentially) and accompanied by large uncertainties. Furthermore, the regression analysis of the effect of treatment on the total H_2_O_2_ dissociation capacity adjusted for AZD-resistant H_2_O_2_ dissociation for the whole small intestine (with intestinal segment introduced as a fixed effect) (not shown) did not provide evidence to confirm the aforementioned hypothesis. In regards to the H_2_O_2_ dissociation rate, the interaction between intestinal segment and treatment was observed as the dissociation capacity was increased in the colons of the intrastriatally treated animals (+76% in 6-OHDA vs CTR; 2.64-fold in CIS vs CTR). The total reductive capacity demonstrated a trend toward increased antioxidant capacity in intrastriatally treated animals in concordance with other redox biomarkers (e.g. reduced TBARS and increased GSH in the duodenum (CIS, 6-OHDA), and increased GSH in the colon (CIS)). Multivariate analysis revealed no evident patterns that would indicate a dysregulation of redox homeostasis in any group (Fig 3).

**Table 1.**
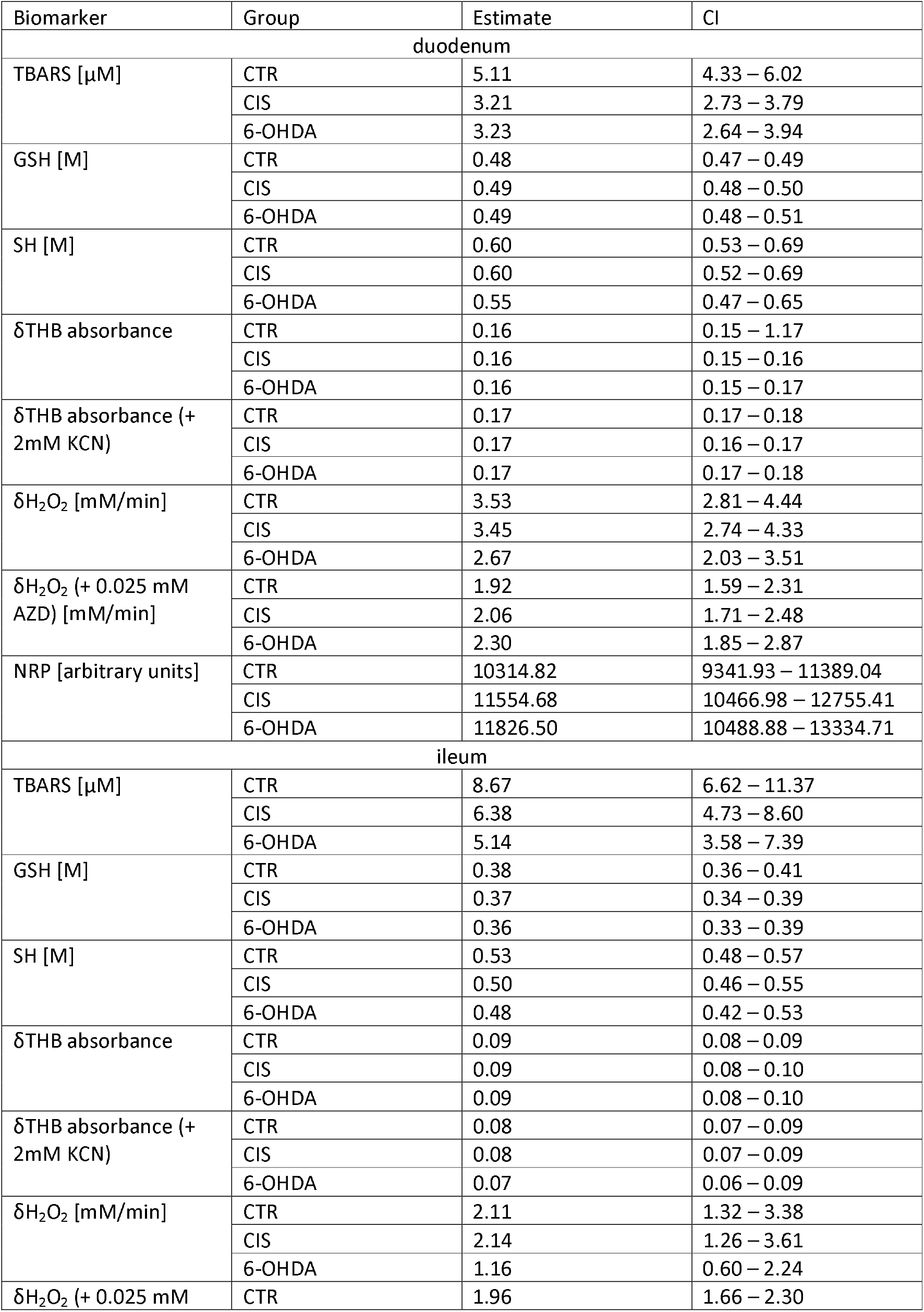
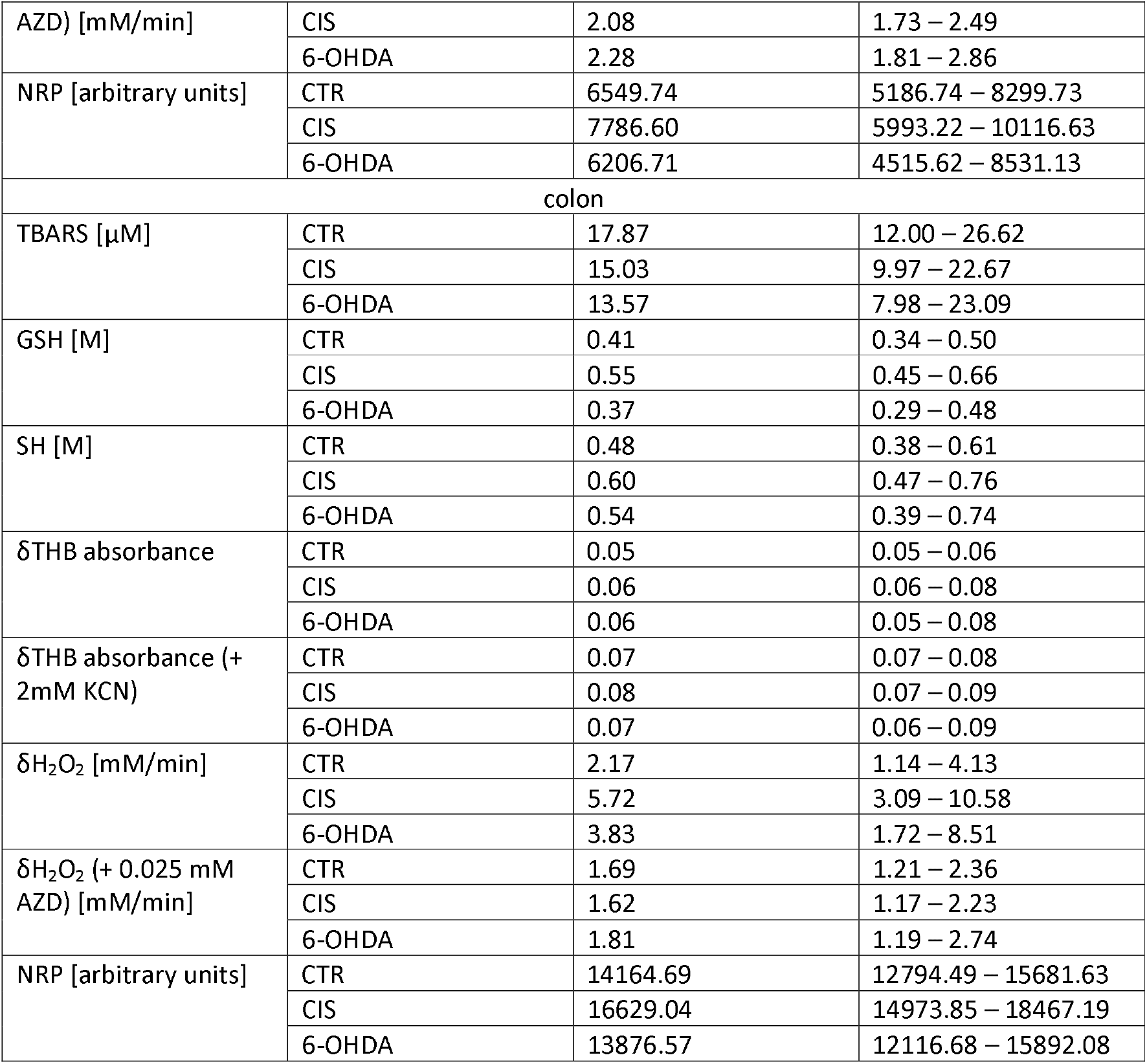
Redox biomarkers in duodenum, ileum, and colon of the control intact (CTR), vehicle-treated animals (CIS), and rats 3 months after bilateral intrastriatal administration of 6-hydroxydopamine (6-OHDA).

**Fig 2.**
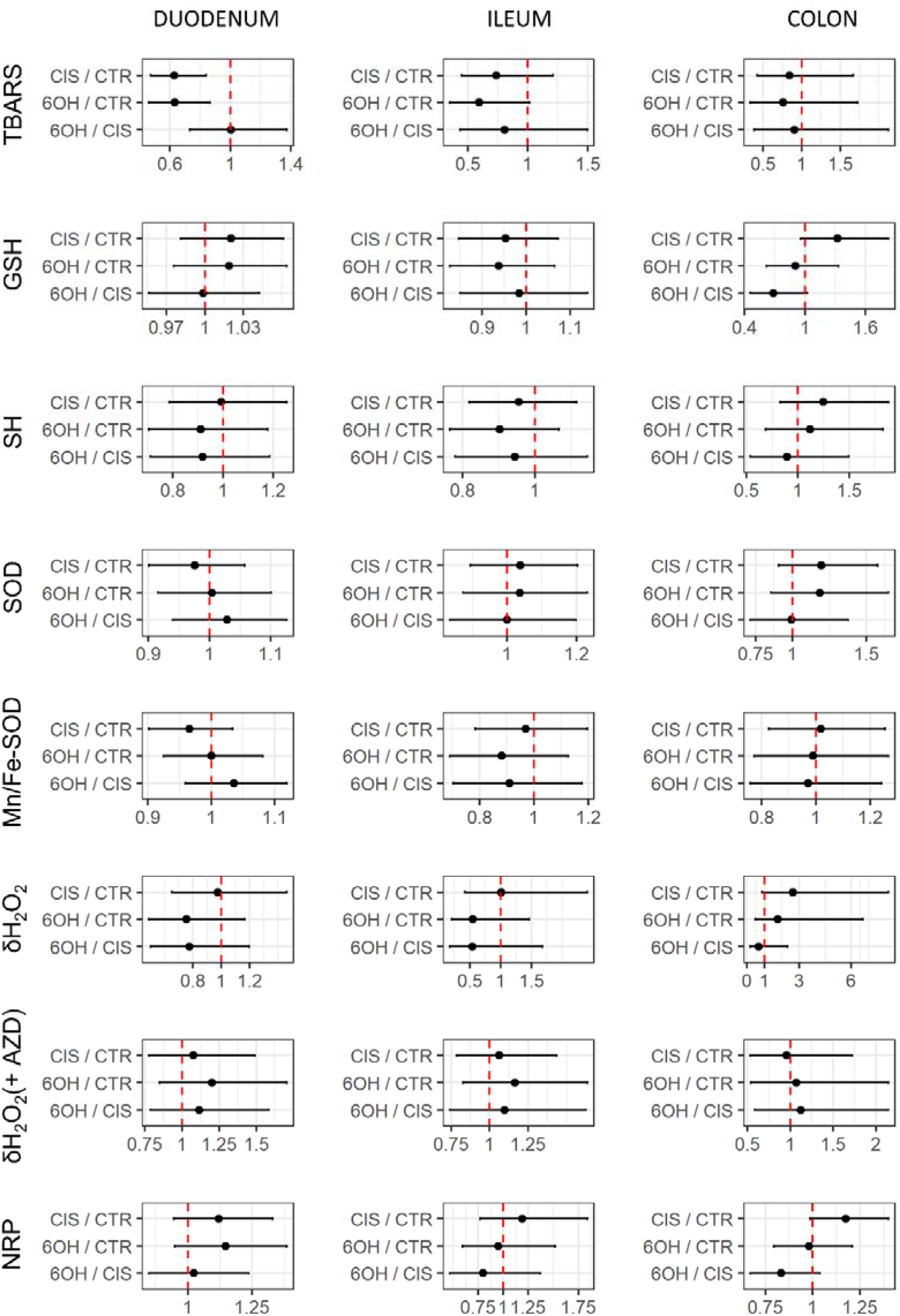
Comparison of redox biomarkers in the control, 6-OHDA, and vehicle-treated animals. Point estimates of ratios of estimated marginal means with respective 95% confidence intervals are shown. CTR – intact controls; CIS – vehicle-treated controls; 6-OHDA – experimental group treated with bilateral intrastriatal 6-hydroxydopamine; TBARS – thiobarbituric acid reactive substances; GSH – glutathione; SH – free sulfhydryls; SOD – superoxide dismutase; AZD – sodium azide; NRP – nitrocellulose redox permanganometry.

**Fig 3.**
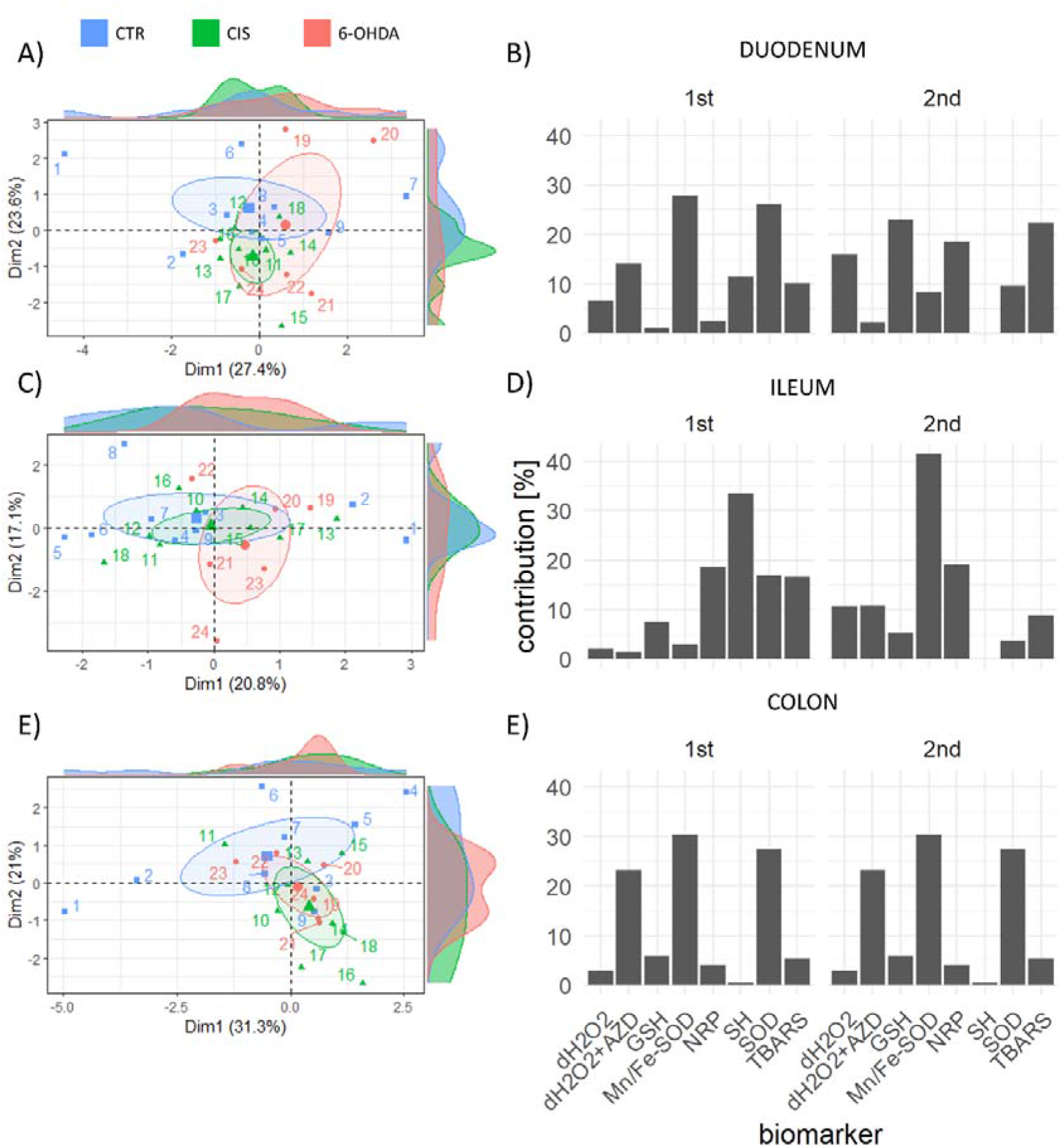
PCA of redox-related biomarkers in the duodenum, ileum, and colon of the brain-first 6-OHDA-induced rat model of PD. A) A biplot of individuals showing the results of multivariate exploration in regards to redox biomarkers in the duodenum. B) Contribution of individual variables to the first two principal components obtained by the PCA of redox-related biomarkers measured in the duodenum. C) A biplot of individuals showing the results of multivariate exploration in regards to redox biomarkers in the ileum. D) Contribution of individual variables to the first two principal components obtained by the PCA of redox-related biomarkers measured in the ileum. E) A biplot of individuals showing the results of multivariate exploration in regards to redox biomarkers in the colon. F) Contribution of individual variables to the first two principal components obtained by the PCA of redox-related biomarkers measured in the colon. PCA – principal component analysis; 6-OHDA – 6-hydroxydopamine; PD – Parkinson’s disease; Dim1 – 1^st^ dimension; Dim2 – 2^nd^ dimension; dH2O2 – the difference in hydrogen peroxide concentration; dH2O2+AZD – the difference in hydrogen peroxide concentration in presence of sodium azide; GSH – glutathione; NRP – nitrocellulose redox permanganometry; SH – free sulfhydryls; SOD – superoxide dismutase activity.

## Discussion

The presented results provide solid evidence against the existence of pronounced gastrointestinal redox dyshomeostasis in a brain-first rat model of PD induced by bilateral intrastriatal administration of 6-OHDA 12 weeks after model induction. Although the possibility that redox dyshomeostasis was present but too subtle to be estimated with a high degree of certainty using the current experimental design and methodological approach cannot be completely excluded, a substantial dropout rate and pronounced motor deficits (Fig 1) accompanied by unconvincing alterations of 8 individual redox biomarkers in 3 separate anatomical regions strongly speak against the presence of gastrointestinal redox dyshomeostasis and/or its biological significance. The latter is further supported by the fact that the same biomarkers have been shown to be sufficiently sensitive to detect redox-related alterations induced by relatively mild stimuli (e.g. single orogastric administration of the 200 mg/kg D-galactose solution)[37] and in other neurotoxin-based models of brain-first-induced neurodegeneration [31, 32].

The data presented here seem to be in contrast with the results reported by Pellegrini et al. who found evidence supporting increased lipid peroxidation in the colon of the 6-OHDA-treated rats 4 and 8 weeks after model induction in the only other study in which oxidative stress-related biomarkers were measured in the gastrointestinal tract of the brain-first 6-OHDA-induced model of PD [48]. Nevertheless, the apparently discrepant results may be explained by several methodological differences. Pellegrini et al. used the 6-OHDA model in which the toxin is injected unilaterally into two sites of the medial forebrain bundle (MFB) in a total dose of 7.5 μg/3 μl [48] and we used bilateral intrastriatal administration of 8 μg/2 μl into each hemisphere to overcome potential compensatory mechanisms of the un-lesioned hemisphere that may mask some of the effects of the nigrostriatal dopaminergic neurodegeneration [49]. Both the dose and the site of injection have a profound effect on the model phenotype and spatiotemporal patterns of dopaminergic neurodegeneration. While administration of 6-OHDA into the SN and/or the MFB induces a complete and rapid lesion of the nigrostriatal pathway in hours, striatal injection usually produces a slow, progressive, and partial damage spreading from the striatal terminals to SN for weeks [50]. Considering that in our experiment SN lesion was minimal even after 12 weeks (assessed by control SN TH immunopositivity [51]), it is possible that the absence of the effects in the gut observed here was due to relative preservation of the nigro-vagal pathway (the main pathway implicated in the gut-related PD symptoms [52]). Furthermore, Pellegrini et al. measured only a single redox biomarker – malondialdehyde (MDA) using the TBARS method [48]. Although TBARS is a widely used and reliable biomarker of oxidative stress [53] its use has many caveats and limitations and it has been proposed that it can offer „at best, a narrow and somewhat empirical window on the complex process of lipid peroxidation“[54]. MDA is one of many different aldehydes that are produced as secondary products of lipid peroxidation and its concentration inside cells depends on many biological processes [53, 55]. Importantly, an increased concentration of MDA can sometimes reflect activation of protective antioxidant defense systems in case its metabolism and the rest of the redox regulatory network work correctly [55]. Considering that MDA is also produced in the process of prostaglandin biosynthesis, especially the synthesis of thromboxanes via the thromboxane synthase [53] implicated in the regulation of inflammation in the gastrointestinal tract [56], it is possible that increased colonic MDA reported by Pellegrini et al. was more related to the observed pro-inflammatory signaling (increased tumor necrosis factor α and interleukin-1β [48]) than a complete failure of the redox homeostasis resulting in uncontrolled peroxidation of membrane lipids. Nevertheless, inflammatory and redox processes demonstrate a high level of biological interdependence [57] so in the context of pro-inflammatory alterations reported by Pellegrini et al. redox dysregulation can by no means be excluded. Finally, Pellegrini et al. measured TBARS in „colonic neuromuscular tissue“, and here we measured redox homeostasis in the intact whole preparations to i) overcome potential bias introduced by dissection and sample preparation; and ii) take into account alterations of the mucosa due to its critical importance for the maintenance of the redox homeostasis in the gastrointestinal tract [58].

## Conclusion

To conclude, measurements of 8 individual redox biomarkers in 3 anatomical regions of the gastrointestinal tract (duodenum, ileum, and colon) indicate that bilateral intrastriatal administration of 6-OHDA can induce pronounced motor deficits without affecting gastrointestinal redox homeostasis. Considering that SN was still not considerably damaged (after 12 weeks after 6-OHDA) the present results suggest that dysfunction of the nigro-vagal system may be a key mediator of gastrointestinal dyshomeostasis in brain-first 6-OHDA-induced rodent models of PD. Consequently, the dose and the site of the 6-OHDA injection may have profound effects on the development of pathophysiological alterations in the gastrointestinal tract and should be carefully considered when brain-first 6-OHDA models are used for studying gastrointestinal symptoms associated with PD.

## Limitations

Finally, several limitations of the present work have to be emphasized. In the present study, there was a pronounced drop-out rate in the 6-OHDA-treated group of animals (57% fatality rate) which may have introduced attrition bias – e.g. the animals with the most pronounced response to the 6-OHDA administration, and possibly more rapid spreading of the 6-OHDA-induced damage from the striatal terminals to SN that may have developed gastrointestinal redox dyshomeostasis were excluded from the study as they died before the final time-point. Furthermore, the main aim of the study was to assess redox homeostasis so additional functional gastrointestinal parameters were not measured. Animal models of PD often demonstrate gastrointestinal dysmotility (hypothesized to be mediated by the degeneration of the nigro-vagal pathway [52] so the association (or the lack of thereof) between gastrointestinal dysmotility and gastrointestinal redox homeostasis remains to be explored in future studies. The latter seems particularly interesting as redox disbalance has been recognized as an important regulator of gastrointestinal motility [59, 60] and it has been hypothesized that motor activity of the gastrointestinal tract may modulate gastrointestinal redox homeostasis [61]. Lastly, gastrointestinal redox homeostasis may be influenced indirectly by behavioral and metabolic consequences of 6-OHDA administration (e.g. altered circadian activity and feeding patterns). Considering that neither circadian activity nor feeding patterns were monitored in the present study and included in statistical models it cannot be excluded that unavoided bias at this level may have masked some of the effects. Future studies should take into account the possible indirect effects of 6-OHDA in this context as well.

## Authors’ contributions

JH, ABP, AK, and JOB – *in vivo* part of the experiment, tissue collection. JH, MJ, GG, ES – biochemical measurements. JH – data curation, data analysis, writing the first draft of the manuscript. MJ, GG, ES, DV, ABP, AK, JOB, and MSP – critical revision of the manuscript. MSP – funding, supervision.

## Data availability statement

Raw data can be obtained from the corresponding author.

## Ethics approval

Only certified researchers worked with the animals and the experiment was conducted with the highest standard of animal welfare. The animal procedures were conducted in concordance with current institutional (University of Zagreb School of Medicine), national (The Animal Protection Act, NN135/2006; NN 47/2011), and international (Directive 2010/63/EU) guidelines governing the use of experimental animals. The experiments were approved by the Croatian Ministry of Agriculture (EP 186 /2018) and the Ethical Committee of the University of Zagreb School of Medicine (380-59-10106-18-111/173).

